# Possible functions of the conserved peptide encoded by the RNA-precursor of miR-156a in plants of the family *Brassicaceae*

**DOI:** 10.1101/2020.10.29.361550

**Authors:** T.N. Erokhina, D.Y. Ryazantsev, L.V. Samochvalova, A.N. Orsa, S.K. Zavriev, S.Y. Morozov

## Abstract

Recent studies have shown that the primary transcripts of some microRNA genes (pri-miRNAs) are able to express short proteins (peptides) ranging usually from 12-15 amino acid residues to around 30 residues in length. These peptides, called miPEPs, may participate in the regulation of transcription of their own pri-miRNAs. Using bioinformatic comparative analysis of pri-miRNA sequences in plant genomes, we previously discovered a new group of miPEPs (miPEP-156a), which is encoded by pri-miR156a in several dozen species from the *Brassicaceae* family. Exogenous peptides miPEP-156a can effectively penetrate plant seedlings through the root system and spread systemically to the leaves of young seedlings. At the same time, a moderate morphological effect is observed, which consists in accelerated growth of the main root of the seedling. In parallel, a positive effect is observed at the level of pri-miR156a expression. It is important that the effects at the morphological and molecular levels are seemingly related to the ability of the peptide to quickly transfer into the cell nuclei and bind to nuclear chromatin. In this work, the secondary structure of the peptide was also experimentally established, and changes in this structure in the complex with DNA were shown.

## 1. Introduction

MicroRNAs (miRNAs) are short, non - coding, endogenous RNAs that regulate gene expression using a specific cleavage and/or translational repression of the target mRNA due to complementary base pairing (Bartel, 2004; Rogers and Chen, 2013; Yu et al., 2017; Wang et al., 2019). Typical miRNAs act as key regulators of various processes of plant and animal development (Liu et al., 2018; Song et al., 2019). Biogenesis of miRNAs requires a number of the well-coordinated processes, where the miRNA gene is first transcribed by RNA polymerase II to form a primary miRNA (pri-miRNA), which usually has a length of several hundred to even thousands of nucleotides, and contains, like mRNA, a 5’-terminal “cap” structure and a 3’-terminal poly(A) sequence (Bartel, 2004; Rogers and Chen, 2013; Yu et al., 2017; Wang et al., 2019; Budak et al., 2017). In plant nuclei, pri-miRNAs are processed to produce a hairpin-like imperfect duplex called a miRNA precursor or pre-miRNA, and this step is performed by Dicer-like enzyme (DCL) involving the HYL1 protein (Fukudome and Fukuhara, 2017). At the next processing step, the loop of hairpin structure of pre-miRNA is cleaved off by DCL, and subsequent 3’-terminal methylation leads to the formation of mature miRNAs (Bartel, 2004; Rogers and Chen, 2013; Yu et al., 2017; Wang et al., 2019; Liu et al., 2018; Song et al., 2019; Budak and Akpinar, 2017; Fukudome and Fukuhara, 2017).

In the cytoplasm, mature miRNA interacts with the RISC protein complex containing the Argonaut enzyme (AGO), which selects one of the chains of double-stranded miRNA after duplex melting. Further complementary base-pairing with mRNA either provides precision cleavage of the target transcript, or inhibits its translation (Bartel, 2004; Rogers and Chen, 2013; Yu et al., 2017; Wang et al., 2019; Zhang et al., 2015). Obviously, the main role of pre-miRNAs is the production of regulatory miRNAs, and there was a common opinion that sequences positioned to the 5’- and 3’-ends from the hairpin-like region of pri-miRNA are functionally unimportant and quickly degrade after cleavage of the pre-miRNA region (Song et al., 2019; Budak and Akpinar, 2017; Fukudome and Fukuhara, 2017).

However, recent methods of plant translatome search suggest the translational activity of pri-miRNA. To facilitate the capture of ribosome-associated RNAs, a method for translational immunological purification of ribosome-bound RNAs (TRAP - translating ribosome affinity immunopurification) was developed (Tavormina et al., 2015). The epitope-labeled ribosomal protein L18 (RPL18) embedded in functional ribosomes in the transgenic *Arabidopsis thaliana* was initially used for affinity purification. The main advantage of this method is that the contamination of polysome preparations with mRNA–protein complexes of similar density is reduced, and these preparations can be used to obtain mRNA fragments directly associated with ribosomes (Tavormina et al., 2015; Zanetti et al., 2005). TRAP analysis has identified polysome preparations containing pri-miRNA fragments associated with ribosomes (Zanetti et al., 2005; Juntawong et al., 2014).

Currently an increasing number of reports indicate that there are indeed short ORFs in the 5 ‘-proximal pri-miRNA sequences coding for peptides. Such peptides encoded by pri-miRNA in plants and animals are called miPEPs (Zhu et al., 2018; Hellens et al., 2016; Li et al., 2017; Matsumoto and Nakayama, 2018; Yeasmin et al., 2018). In particular, a number of miPEPs, whose synthesis is directed by short ORFs in the 5’-proximal pri-miRNA sequences, was found in Arabidopsis (*Arabidopsis thaliana*) (Lauressergues et al., 2015), soy (*Glycine max*) (Couzigou et al., 2016), and peanuts (*Arachis hypogaea*) (Ram et al., 2019). External treatment of plants with synthetic peptides has shown that the peptides miPEP165a from Arabidopsis and miPEP171b from *Medicago truncatula* are able to activate transcription of their own pri-miRNA. Thus, a positive feedback loop is formed, which leads to an increase in the level of biogenesis of the corresponding microRNA (Lauressergues et al., 2015; Couzigou et al., 2015, 2016). Treatment of *M. truncatula* plants with the synthetic peptide miPEP171b increases endogenous miR171b expression, which leads to a decrease in lateral root density. This effect of miPEP171b was specific because the peptide did not affect the expression of other microRNAs. Treatment of *A. thaliana* seedlings with miPEP165a also resulted in a specific increase in miR165a accumulation (Lauressergues et al., 2015; Couzigou et al., 2015, 2016). Recently, the participation of miPEP172c in the control of nodulation in soybean plants was shown (Couzigou et al., 2016).

Previously, we predicted the existence of the short conserved ORFs in the 5’-proximal sequences of pri-miR156a RNA transcripts in plants of the genus *Brassica* (*B. napus*, *B. rapa*, and *B. oleracea*). Similar short conserved peptides with a length of 32-33 residues are also encoded in the genomes of several plants from the genus *Arabidopsis*, including *A. thaliana* (Morozov et al., 2019). Using multiple alignments of the coding sequences of the miPEP-156a micropeptides, conservative amino acid residues were identified in many representatives of 11 genera in the *Brassicaceae* family (Morozov et al., 2019). The secondary structure and three-dimensional model of miPEP-156a were predicted *in silico* using the I-TASSER (Iterative Threading ASSEmbly Refinement) method. It was found that the secondary structure of all peptides contains α-helices, and miPEP-156a in the genus *Brassica* is a potential dimeric or oligomeric protein (Morozov et al., 2019). In this work, the secondary structure of the peptide was experimentally established using the synthetic peptide miPEP-156a from *Brassica rapa*. Moreover, the biological and a number of biochemical properties of this peptide were studied.

## 2. Results and discussion

### 2.1. Comparative analysis of accumulation of pri-miR156a transcripts in plants of genera Brassica and Arabidopsis

Previously, it was shown that the most intensive synthesis of pri-miR156a in the leaves and roots of Arabidopsis seedlings occurs during germination between the 2nd and 10th day at 23°C (Kim et al., 2016; Yu et al., 2015). After the 10th day, this expression drops significantly. We decided to compare the dynamics of pri-miR156a accumulation in broccoli and Chinese cabbage plants (*Brassica oleracea* var. italica and *Brassica rapa* subsp. pekinensis) in comparison with Arabidopsis plants. For this purpose, total RNA was isolated from the leaves and roots of 3-4-day-old seedlings, as well as flowering plants, and quantitative PCR analysis was performed using a cDNA preparation obtained using oligo(dT) primers (Fig. 1). Importantly, such comparative studies for cabbage plants were not performed previously.

**Fig. 1.**
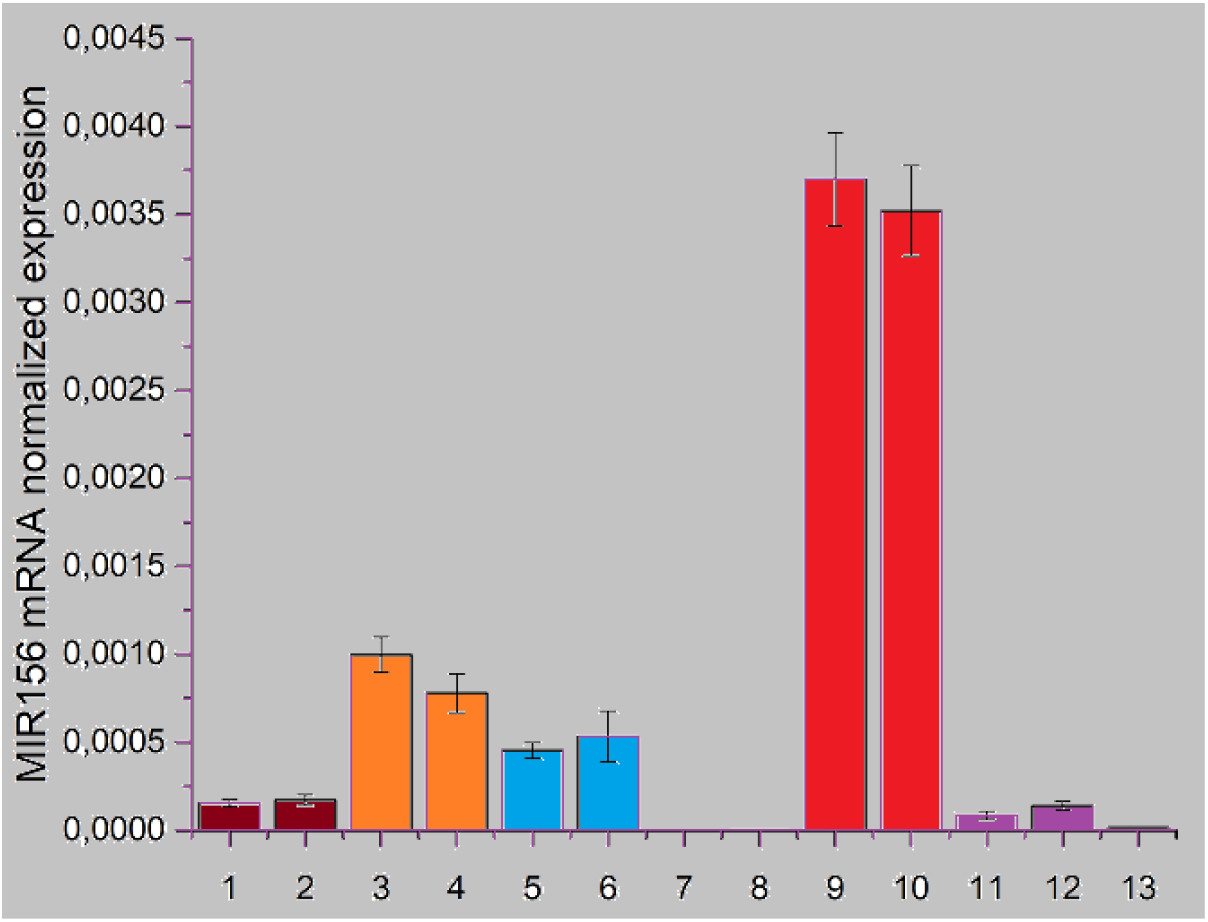
Quantitative PCR for measuring the level of pri-miRNA156a expression in plants of the *Brassicaceae* family. (1) Broccoli (20-day plant, leaves), (2) broccoli (20-day plant, root), (3) broccoli seedlings (4-day plant) with miPEP156a peptide added (10 μg/ml), (4) broccoli seedlings (4-day plants) with peptide added (10 μg/ml), (5) broccoli seedlings (4-day plants), control (water), (6) broccoli seedlings (4-day plants), control (water), (7) adult flowering Arabidopsis plants (roots), (8) adult flowering Arabidopsis plants (leaves), (9) Arabidopsis, whole seedlings (4-day plants), (10) Arabidopsis, whole seedlings (4-day plants), (11) Chinese cabbage (20-day plant, stem), (12) Chinese cabbage (20-day plant, leaf), (13) Chinese cabbage (20-day plant, root).

It was found that the ontogenetic correlation of plant age and the accumulation of pri-miR156a in leaves and roots of plants is quite similar in cabbage and Arabidopsis. The accumulation of pri-miR156a in seedlings is many times higher than in flowering plants (Fig. 1). Therefore, the works on the activities and physical identification of miPEP156a in plants should be carried out at the early stages of ontogenesis.

### 2.2. Bioinformatic analysis of Arabidopsis translatome

Previously, researchers raised the question of which approaches can be used to test the expression of potential miPEPs in plants (Lv et al., 2016). For example, miPEP165a of Arabidopsis was not detected by mass spectrometry (Lauressergues et al., 2015), although TRAP analysis clearly showed that pri-miRNA fragments encoding this peptide can be detected in polysome preparations (Juntawong et al., 2014). In our preliminary experiments, attempts to detect miPEP-156a in Arabidopsis seedlings using immunological approach and mass spectrometry also failed (data not shown). Thus, in order to obtain more direct evidence of the coding ability of the 5’-terminal ORF in pri-miR156a in Arabidopsis plants, we performed a bioinformatic BLASTn analysis of sequences in the translatome based on the NCBI SRA archive (SRA – short read assembly).

The Arabidopsis pri-miR156a (AT2G25095) gene is located on chromosome 2 and has a size of 3108 base pairs. This gene consists of 4 exons and 3 introns. The pre-miR156a hairpin sequence is located in exon 2, and the miPEP165a ORF is located in exon 1. For the BLASTn analysis, the query sequence of miPEP165a ORF region and the similarly sized region from exon 4 were taken. The analysis of data from a number of TRAP experiments revealed the presence of miPEP156a ORF in ribosome-bound RNA fragments from the roots of seedlings [NCBI accessions SRX3204187, SRX3204194, SRX3204195, SRX3204199 (Brosnan et al., 2019)], although sequences of the 3’-terminal exon 4 were not found among ribosome-bound segments (data not shown). Thus, we obtained direct evidence that the 5’-terminal ORF on the pri-miR156 RNA may undergo translation, at least in the roots of *A. thaliana* seedlings.

### 2.3. Effect of exogenous miPEP156a peptide on root development in plant seedlings

Since Arabidopsis miPEP165a is known to modulate the accumulation of mature microRNA by enhancing the transcription of its precursor RNA transcript and therefore affects the rate of development of seedling roots (Lauressergues et al., 2015), we assumed that the application of exogenous miPEP156a can also affect the development of the root system of young cabbage seedlings. To test whether miPEP156a is physiologically active, Chinese cabbage seeds (*Brassica rapa* subsp. pekinensis) were germinated on a plant culture medium supplemented with a synthetic peptide at concentration 10.0 μg/ml. It was found that the length of the main (primary) root of 5-day-old seedlings grown on a medium supplemented with exogenous miPEP156a increased compared to the control (water solution of salts) (Fig. 2). This increase in root length is comparable to the effect described in the literature for Arabidopsis seedlings overexpressing miR156a, where seedlings with increased miR156a expression had longer primary roots (about 20%) than the wild type (Yu et al., 2015). On the contrary, under conditions of suppression of miR156a synthesis, shorter primary roots developed in comparison with the wild type plants (Yu et al., 2015).

**Fig. 2.**
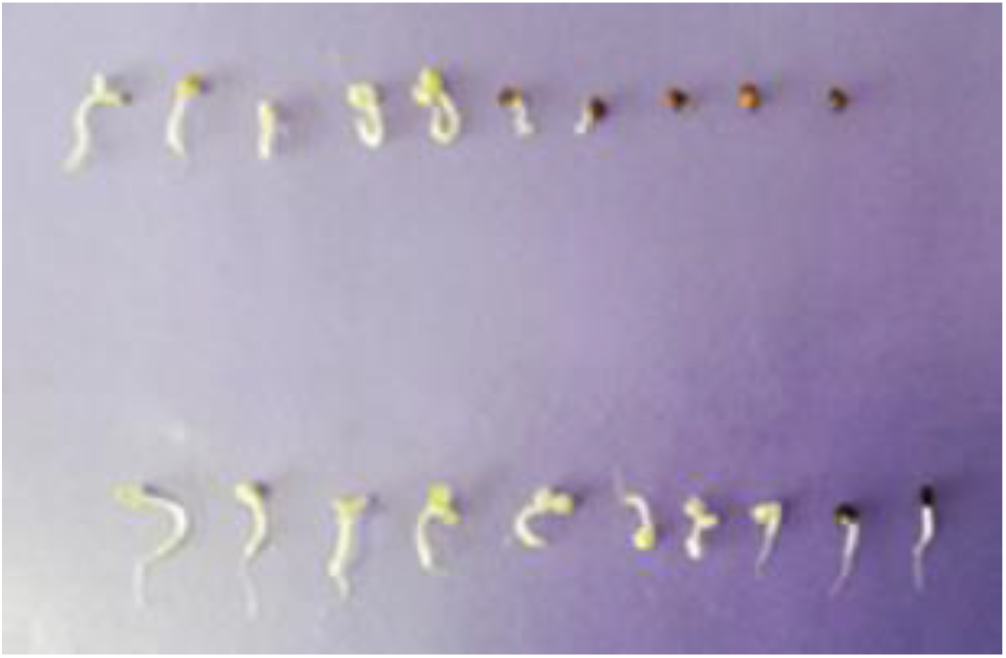
Germination of cabbage seeds in the presence and absence of miPEP156a. Top row – sprouting without miPEP156a (average root length for 50 plants – 0.7 + 0.7 cm); bottom row - sprouting with the addition of miPEP156a (concentration of 10 μg/ml) (average root length for 50 plants – 1.6 + 0.9 cm).

These data allow us to hypothesize that miPEP156a plays a role in increasing miR156 accumulation and indirectly in the formation of primary roots in cabbage. This is very similar to the effect of miPEP165a of Arabidopsis on the growth of primary roots and increased accumulation of the corresponding microRNA (Lauressergues et al., 2015). Indeed, direct measurement of the amount of pri-miRNA156a in broccoli sprouts grown in the presence of the peptide revealed a clear (although rather moderate) increase in the concentration of this RNA in whole plants (leaves, stem, roots) (Fig. 1).

### 2.4. Localization of exogenous miPEP156a peptide in plant seedlings

Finding of the physiological effect of exogenous miPEP156a on plant growth indicates its ability to penetrate the tissues of seedlings. Previously, this effect was studied in detail on the example of miPEP165a in *A. thaliana* (Ormancey et al., 2020). This published study used a peptide labeled with a fluorescent dye. It was shown that in 7-day-old Arabidopsis seedlings, the labeled peptide actively penetrates the root epidermis and root meristem during 2 hours of incubation. However, it takes longer time to penetrate other parts of the root. It was found that two mechanisms are involved: primary passive diffusion of the peptide followed by active endocytosis (Ormancey et al., 2020). It is important that during these processes, miPEP165a cannot penetrate the central cylinder of the root, so it does not enter the phloem. It is correlated with the fact that peptide does not migrate from the roots to the upper part of the plant (Ormancey et al., 2020). Our experiments with the penetration of miPEP156a into cabbage seedlings revealed a different picture. Incubation with TAMRA-miPEP156a (Table 1) showed that only small amounts of the fluorescent peptide are localized in the roots of seedlings after 18 hours of incubation. In this case, only the nuclei of elongated cells in the central cylinder of the root (presumably phloem parenchyma cells and companion cells) are strongly labeled (Fig. 3A). Microscopic analysis showed that, in contrast to miPEP165a, the most intense accumulation of TAMRA-miPEP156a is observed in the seedling leaves. Moreover, the accumulation of the peptide is clearly higher in the leaf areas near the veins, which suggests the miPEP156a transport through the phloem from the roots to the upper part of the seedling. In leaf cells, where a sufficient amount of the peptide is found, most labelling is observed near and inside the nucleus (Fig. 3B).

**Table 1.**
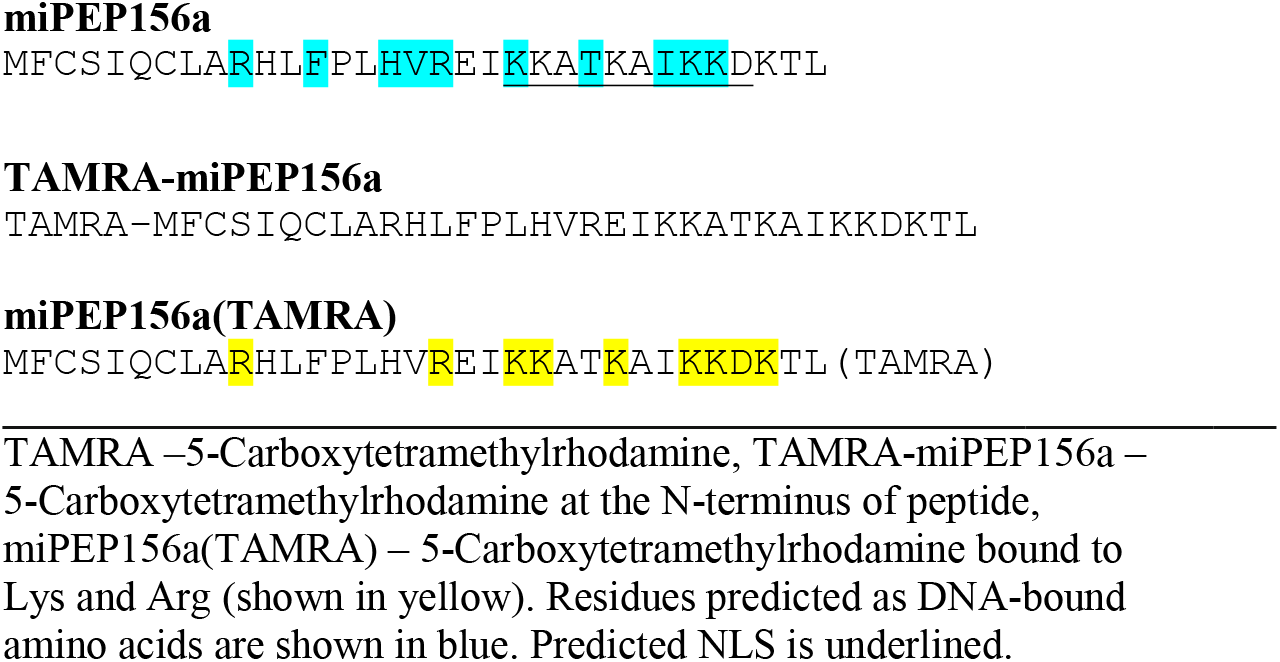
Peptides used in this study

**Fig. 3A.**
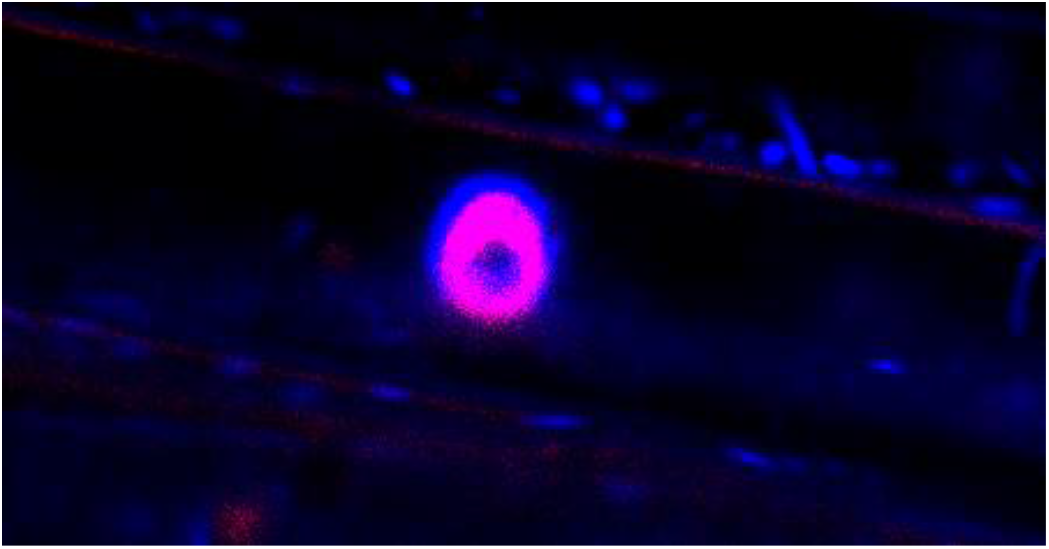
Confocal microscopy of the root of a cabbage seedling incubated in the presence of the fluorescent peptide TAMRA-miPEP156a (red). Hoechst 33342 nucleus-specific dye is in blue.

**Fig. 3B.**
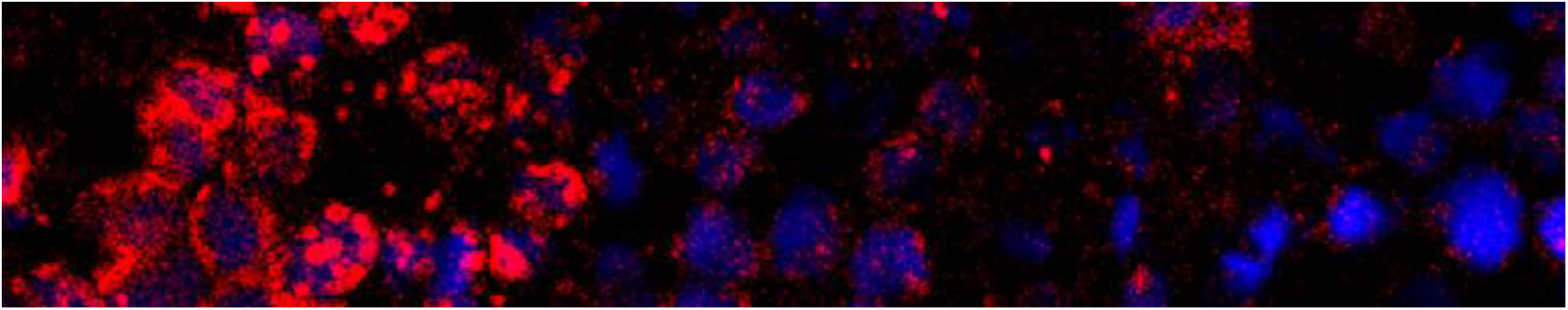
Confocal microscopy of cabbage leaf when seedling was incubated in the presence of the fluorescent peptide TAMRA-miPEP156a (red). Hoechst 33342 nucleus-specific dye is in blue. The left part of the image corresponds to the part of the leaf localized near the central vein.

### 2.5. Study of miPEP156a peptide localization in animal cells

The observed localization of miPEP156a in the nuclei of cabbage plant cells, which was not detected in the case of miPEP165a in *A. thaliana* (Ormancey et al., 2020), indicates the ability of miPEP156a peptide to efficient nuclear-cytoplasmic transport. The mechanisms of such transport are very conservative in plants and animals (Sorokin et al., 2007; Merkle, 2011). Therefore, we selected Sp2/0 mouse myeloma cell culture as the target for monitoring the effectiveness of nuclear import of the miPEP156a peptide. The cells were incubated with TAMRA-miPEP156a for 2 hours, washed, and analyzed using a confocal microscope.

Preliminary staining of Sp2/0 cells with the Hoechst 33342 nuclear marker and fluorophore staining the endoplasmic reticulum showed that the cell nucleus occupies a large part of the cell volume, and there is only a thin layer of cytoplasm adjacent to the plasma membrane (Fig. 4A and B). After visualizing the cells double stained with the miPEP156a and Hoechst 33342, it has become obvious that the fluorescent peptide is almost completely imported into the nucleus and hardly observed in the cell cytoplasm (Fig. 5A and B). These data support the active transport of the peptide into the nucleus and cannot be explained solely by its passive diffusion through the nuclear pores.

**Fig. 4A.**
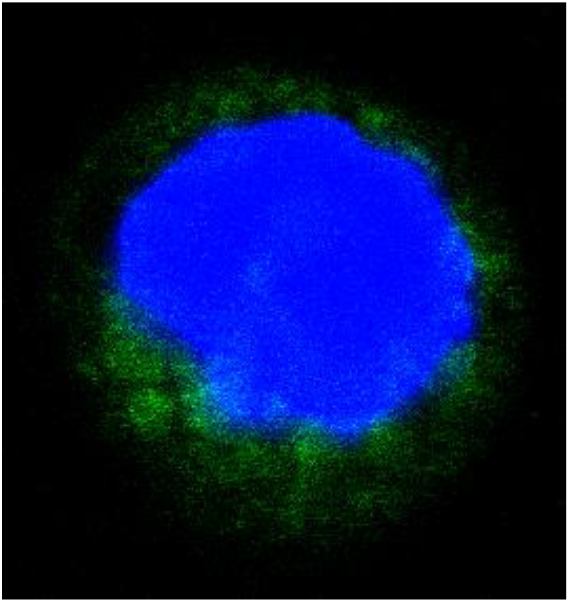
Confocal microscopy of a mouse myeloma cell Sp2/0. Hoechst 33342 nucleus-specific dye is in blue, and cytoplasmic marker ER-Tracker ™ Green is in green.

**Fig. 4B.**
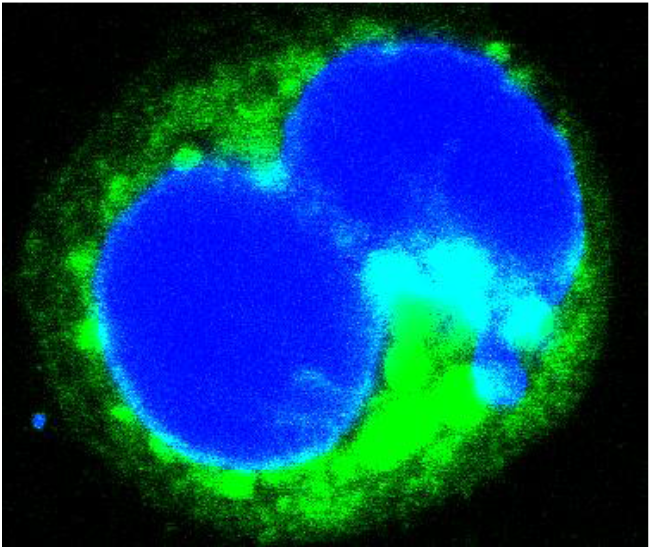
Confocal microscopy of mouse myeloma dividing cell Sp2/0. Hoechst 33342 nucleus-specific dye is in blue, and cytoplasmic marker ER-Tracker ™ Green is in green.

**Fig. 5A.**
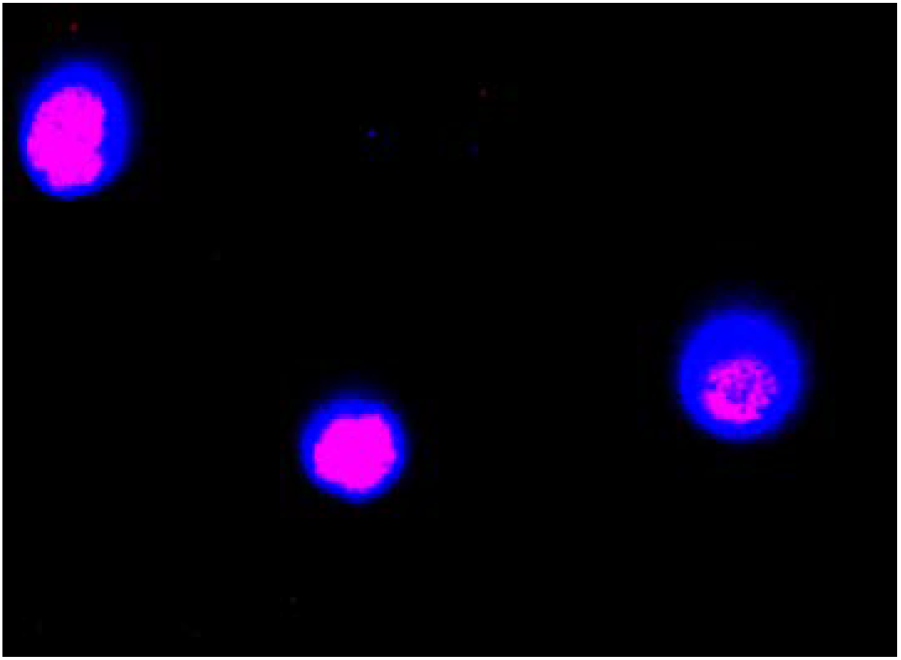
Confocal microscopy of a mouse myeloma cells Sp2/0 incubated in the presence of the fluorescent peptide TAMRA-miPEP156a (red). Hoechst 33342 nucleus-specific dye is in blue. Note that nuclei are only visible.

**Fig. 5B.**
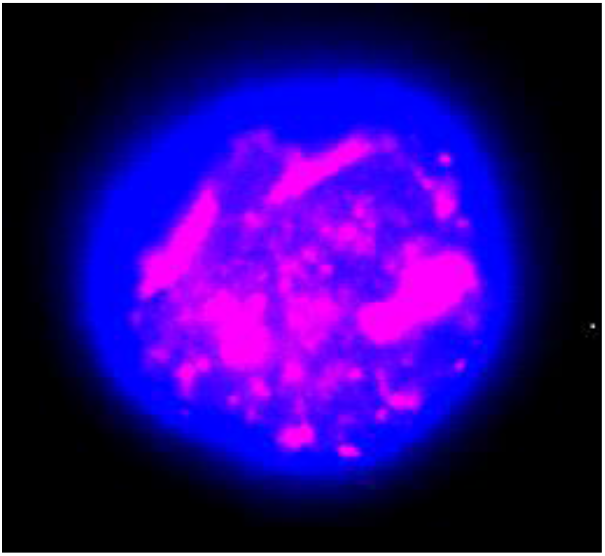
Confocal microscopy at high magnification of a mouse myeloma cell Sp2/0 incubated in the presence of the fluorescent peptide TAMRA-miPEP156a (red). Hoechst 33342 nucleus-specific dye is in blue. Note that nucleus is only visible.

Indeed, the results of an *in silico* analysis of the primary structure of miPEP156a allowed us to hypothesize that the peptide sequence carries a positively charged KKATKAIKKDK segment that includes a nuclear localization signal (NLS) (http://mleg.cse.sc.edu/seqNLS/index.html) (Sorokin et al., 2007; Merkle, 2011) (Table 1). In particular, this segment contains a segment (27)IKKDK(31), quite similar to the KKKGK sequence, which is responsible for the nuclear localization of the fructose 1,6-bisphosphatase enzyme in animal cells. The published results showed that integrity of this motif is crucial for the nuclear localization of the enzyme (Gizak et al., 2009).

### 2.6. In vitro *binding of the miPEP156a peptide to nuclear DNA and plant chromatin*

Localization of miPEP156a in the nuclei of animal and plant cells may indicate its efficient binding to chromatin due to protein-protein and/or DNA-protein interactions. Indeed, the use of computer predictions based on algorithms available at (https://dnabind.szialab.org/) and (http://mleg.cse.sc.edu/DNABind/) (Liu and Hu, 2013) showed that the peptide potentially interacts with DNA with a probability of about 99%, and up to 10 amino acid residues are potentially involved in this process (Table 1). To test the predictions experimentally, we used *in vitro* binding of a fluorescently labeled peptide miPEP156a to cabbage chromatin (Figure 6A). The “building block” of chromatin, the nucleosome, contains ~150 base pairs of DNA wrapped around a histone octamer consisting of four main histones H2A, H2B, H3, and H4 (Lanctot et al., 2007). It has long been established that changes in the chromatin structure required for effective binding of many transcription factors and activation of gene transcription often require removal of H2A–H2B dimers from the nucleosome. Moreover, as a rule, when nucleosomal re-modeling occurs, the H2A-H2B dimer is more easily displaced than H3 and H4 (Bennetzen et al., 2010). *In vivo*, the displacement of the H2A-H2B dimer from the promoter region is usually performed by special nuclear transcription factors (pioneer transcription factors), which often structurally have a three – dimensional structure similar to the H2A-H2B dimer (Lai et al., 2018).

**Figure 6A.**
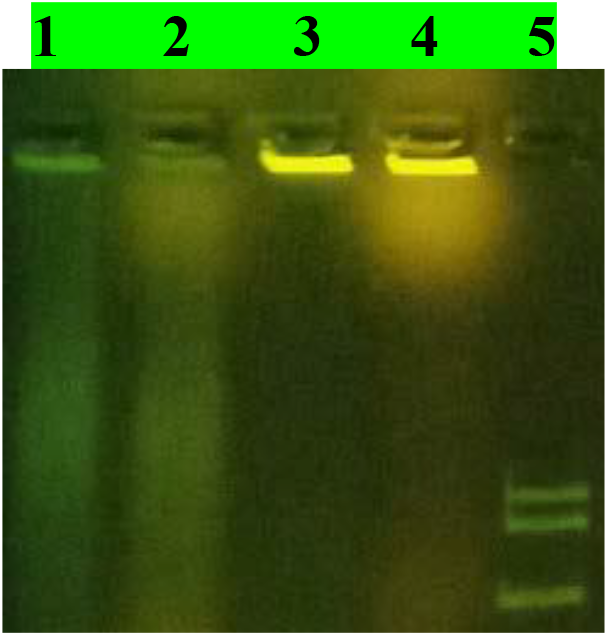
Detection of peptide-chromatin complexes in gel retardation experiment (0.8% agarose gel) after binding of fluorescent peptide TAMRA-miPEP156a with cabbage chromatin. 1 – native chromatin with TAMRA-miPEP156a; 2 - native chromatin with miPEP156a (TAMRA); 3 - temperature-treated chromatin (74°C) with TAMRA-miPEP156a; 4 - temperature-treated chromatin (74°C) with miPEP156a (TAMRA); 5 – size-marker DNA fragments (10000bp, 8000bp, 5000bp).

It can be assumed that if miPEP156a is functionally similar to miPEP165a, as evidenced by our data, the former peptide could interact with chromatin like common transcription factors. Taking into account that some *in vitro* data indicate a direct competition for binding to promoters between transcription factors and the H2A-H2B dimer (Lone et al., 2013), we used *in vitro* two types of cabbage chromatin preparations – the original (“native chromatin”) and chromatin that was subjected to temperature treatment at 74^°^C. It is known that this treatment leads to the displacement of the H2A-H2B dimer, but retains H3 and H4 in the nucleosomes (Horikoshi et al., 2019). It should be noted that the binding efficiency of miPEP156a with chromatin increased for many times after temperature treatment (Fig. 6A). Thus, these results indirectly support the assumption that miPEP156a, like miPEP165a of *A. thaliana*, can modulate the transcriptional activity of genes by interacting with plant chromatin.

In addition, we tested the ability of miPEP156a to interact with isolated plant DNA. The use of the motility shift assay (Rakitina et al., 2011) in agarose gel showed that the peptide has the ability to bind not only the chromatin complex, but also total plant DNA and a non-specific DNA fragment. At high concentrations, the peptide apparently covers a significant part of the length of the DNA molecules, since the mobility of the peptide-DNA complex is significantly reduced (Fig. 6B and C).

**Fig. 6B.**
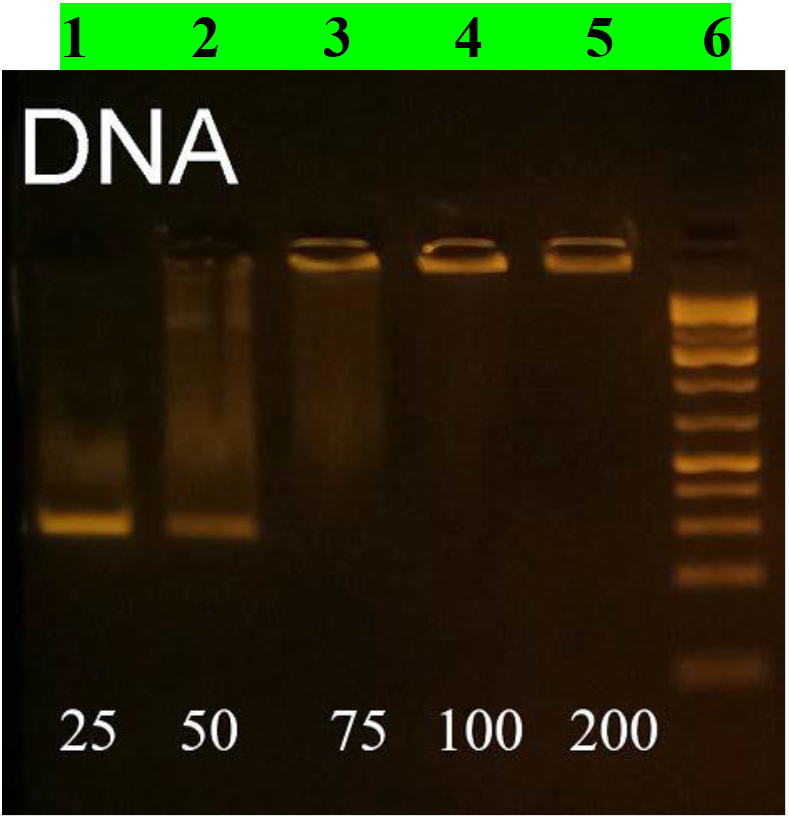
Detection of peptide-DNA complexes in gel retardation experiment (1.5% agarose gel) after binding of fluorescent peptide TAMRA-miPEP156a with a non-specific DNA fragment. Peptide-DNA molar ratio: (line 1) - 25:1, (line 2) - 50:1, (line 3) - 75:1, (line 4) - 100:1, (line 5) - 200:1. Line 6 shows size-marker DNA fragments (from bottom to top 100, 200, 300, 400, 500 bp).

**Fig. 6C.**
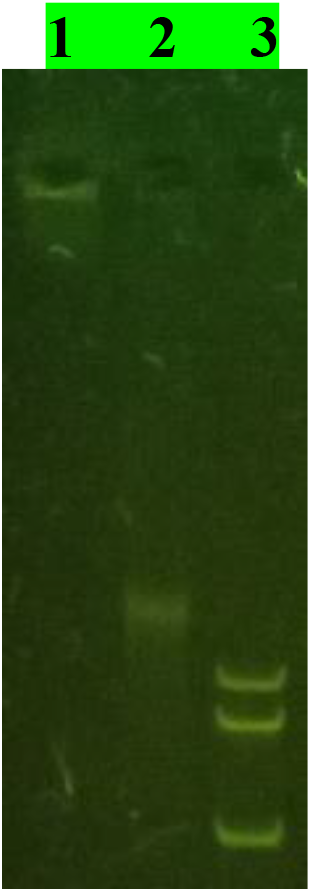
Detection of peptide-chromatin complexes in gel retardation experiment (1.5% agarose gel) after binding of fluorescent peptide TAMRA-miPEP156a with cabbage DNA. 1 – cabbage DNA with TAMRA-miPEP156a; 2 – cabbage nuclear DNA; 3 – size-marker DNA fragments (10000bp, 8000bp, 5000bp).

### 2.7. Secondary structure of the miPEP156a and its complexes with DNA

Among the proteins involved in transcription regulation, a significant proportion of transcription factors interact with chromatin by recognizing DNA sequences through both highly ordered DNA-binding domains and internally disordered regions (IDR - intrinsically disordered regions). IDRs are protein domains that do not have a stable three-dimensional structure under physiological conditions (van der Lee et al., 2014). However, many IDRs adopt a well-defined conformation when interacting with target molecules. A significant number of these proteins interact with chromatin (Peng et al., 2015; Wang et al., 2016). Recently, the interaction of a DNA-binding peptide of the transcription factor GCN4 with model DNA was studied (Quirolo et al., 2019). Importantly, a monomeric peptide, based on GCN4 sequence and including 34-amino acid residues, was not able to bind to a specific dsDNA. Only its dimer, formed through cysteine residues, retains the ability to recognize consensus DNA sequences, which obviously indicates that dimerization is an essential prerequisite for the formation of a peptide-DNA complex. For the GCN4 peptide, it was shown that when bound to DNA, disordered elements can undergo structural transitions that lead to the formation of a well-defined conformation that is stable in the DNA bound state (Miskei et al., 2017).

Interestingly, miPEP156a, which has a length of 33 residues and contains two cysteines, potentially is capable of forming dimers and tetramers (Morozov et al., 2019). Taking into account the formal similarity of the GCN4 peptide and miPEP156a, we experimentally studied the secondary structure of the miPEP156a in the free state and in complex with cabbage DNA. Six independent measurements were performed with two repetitions each at the peptide concentrations 0.012 mM, 0.017 mM and 0.07 mM (Table 2) (Fig. 7). There is an inversely proportional relationship between the number of beta structures and the intensity of the negative signal at 200 nm in the circular dichroism spectrum of miPEP156a (Fig. 7). In the case of GCN4 peptide, which is strongly disordered in a free state, its folding complex with DNA resulted in more alpha helices (Quirolo et al., 2019). However, miPEP156a has an increase in the number of beta structures when it interacts with DNA (Table 2). It is important that the percentage of disordered regions in both peptides significantly decreases after the formation of a complex with DNA.

**Table 2.** Circular dichroism (CD) spectrum of miPEP156a and peptide-DNA complex

| Probe | Alpha helix, % | Beta helix, % | Turn, % | Disordered elements, % | NRMSD |
| --- | --- | --- | --- | --- | --- |
| 1 | $6,9 \pm 0,5$ | $28,5 \pm 0,9$ | $22,4 \pm 0,5$ | $41,2 \pm 0,8$ | 0,043 |
| 2 | $5,5 \pm 0,9$ | $37,4 \pm 2,2$ | $23,0 \pm 0,7$ | $33,9 \pm 1,9$ | 0,06 |

1 – free peptide, 2 – peptide-DNA complex.

**Fig. 7.**
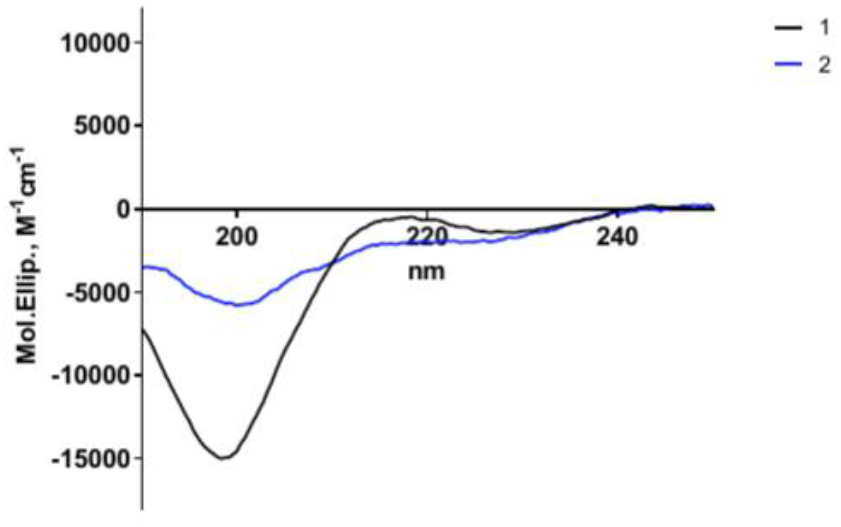
Representative CD spectrum of miPEP156a (1) and its complex with cabbage DNA (2).

## 3. Conclusions

To date, in addition to the miPEP156a found in the *Brassicaceae* family, there are data on several other miPEPs in plants. These peptides are miPEP165a and miPEP858 from *Arabidopsis thaliana* (Lauressergues et al., 2015; Sharma et al., 2020), miPEP171b from *Medicago truncatula* and miPEP172c from *Glycine max* (Couzigou et al., 2015, 2016), and vvi-miPEP171d1 from *Vitis vinifera* (Chen et al., 2020). In our work, we have identified some common features of miPEP156a functioning with the peptides listed above. First, all peptides are able to change plant phenotype upon exogenous application to plant seedlings which represents the overall positive effect on the growth of primary roots. Second, treatment with synthetic peptides shows that miPEPs are able to activate the transcription of their own pri-miRNAs. However, we also observed new effects that were not described in previously published studies. In particular, the penetration of the peptide into the roots of seedlings leads in the case of miPEP156a to its effective transport to the leaves. However, in the case of miPEP165a of *A. thaliana*, the complete retention of the peptide in the roots was observed. The miPEP156a peptide was found only in individual cells of the root phloem. The ability of miPEPs to interact with DNA and transport to the nucleus remained completely outside the scope in published works. Thus, it should be noted that our research has revealed a new spectrum of activities of miPEPs, which is largely consistent with their supposed functions in transcriptional regulation.

## 4. Experimental

### 4.1. Quantitative PCR analysis

To reveal the stages of plant development when the highest expression of the miPEP-156a peptide occurs, experiments on the expression level of pri-miR156a in broccoli cabbage (*Brassica oleracea* var. italica) and Chinese cabbage (*B. rapa* subsp. pekinensis) were conducted. The seeds were sown on fertilized soil and kept at 4^°^C for 3 days before being transferred to the growth chamber (0 day). The plants were grown at a temperature of 22^°^C under fluorescent lamps in a long day (16 hours light/8 hours dark). Plant tissues (3-4 day old whole seedlings) were blended in a liquid nitrogen and total RNA was extracted using TRIzol™ reagent (Invitrogen™, USA) in accordance with the manufacturer’s instructions. After determining the concentration, 2 μg of RNA were incubated with DNAase I (Thermo Fisher Scientific, USA), and, after confirming the quality of RNA by agarose gel electrophoresis with ethidium bromide, reverse transcription was performed with d(T)18-primer using the Reverse Aid H Minus First Strand cDNA Synthesis Kit (Thermo Fisher Scientific, USA). The resulting cDNA was added to the mixture for quantitative PCR, which was performed using reagents from Evrogen (Russia). PEX4 (locus AT5G25760, PEROXIN4 = UBIQUITIN-CONJUGATING ENZYME 21) was used as a reference gene (Czechowski et al., 2005).

For quantitative PCR of the genes encoding MIR156a and PEX4 (control), primers and hydrolyzable probes were selected in such a way that it was possible to amplify the RNA of *A. thaliana*, Chinese cabbage and broccoli. The following set of primers was used: MIR156r d(5’ - CTTTCTTTATGGCTCTTGTCGCTT-3’), corresponding to the conservative sequence at the 3’ end of the miPEP156a ORF in the genera *Brassica* and *Arabidopsis*, MIR156t d(CAGACATCTGTCCCTTTGCATGTAAGA) containing the FAM label and MIR156f d (AAATGTTCTGTTCAATTCAATGC), which corresponds to the 5’-end sequence of this ORF. Quantitative PCR was performed using the mixture qPCRmix-HS (Evrogene, Russia) on the DT prime amplifier (DNA technologies, Russia). The reaction mixture (25 μl) contained 10 pmol of each primer and 5 pmol of the probe. The amplification program was the same for pri-miR156a and PEX4: 95^°^C - 5 min; 95^°^C - 15 sec, 60^°^C - 15 sec*, 72^°^C - 15 sec, in total 45 cycles (* - detection of fluorescence in the FAM channel). After data processing, the Cq values were used to calculate the normalized expression in the QGene program (Simon, 2003).

### 4.2. Study of the effect of the miPEP156a peptide on the development of the root system of plant seedlings

Chinese cabbage seeds were sterilized in a 50% solution of calcium hypochlorite with 0.5% Tween-20 added for 10 minutes while stirring, and then washed six times with sterile distilled water. The seeds were sprouted and grown in a sterile MS medium (Serva, Germany) with the addition of 30 g/l sucrose. For germination, the seeds were placed in flasks (500 ml) with a sterile MS medium and germinated at 80 rpm under a lamp for 24 hours. Next, the sprouted seeds were transferred to sterile 6-well tablets with a sterile MS medium, with or without the addition of miPEP156a peptide at a concentration of 10 μg/ml. For root measure, 4-5-day-old seedlings were transferred to glass plates for observation, measurement, and photography.

### 4.3. Preparation of fluorescently labeled peptides

Synthetic peptides (Table 1) were kindly provided by Igor Ivanov (Shemyakin-Ovchinnikov Institute, RAS). Then we synthesized the fluorescently labeled miPEP156a peptide. TAMRA (5-carboxytetramethylrhodamine) fluorophore was used for tagging. As a result, the TAMRA-miPEP156a peptide was obtained. Peptide miPEP156a was labeled in two different ways: by the N-terminal and by the amino group labelling. To label the peptide by amino groups, 2 mg of the peptide was dissolved in 0.1 M sodium bicarbonate, pH 8.2, and DMSO was added to a 1.5-fold molar excess of N-hydroxysuccinimide ether TAMRA (Lumiprobe, Russia). The reaction was performed for 2 hours at room temperature. After the reaction, the peptide was precipitated with trichloroacetic acid, and the precipitate was washed with acetone and dissolved in water. Reverse-phase chromatography was performed on a Vydac C18 peptide column (Hichrom, UK) in an acetonitrile gradient with the addition of 0.1% trifluoroacetic acid to remove the unreacted dye.

The N-terminal tagging of the peptide was performed in accordance with a previous paper (Witus and Francis, 2010). Pyridoxal phosphate (Sigma, USA) was added to 2 mg of the peptide dissolved in 200 μl of phosphate buffer, pH 6, to a concentration of 10 mM, and incubated overnight at room temperature. After incubation, the mixture was centrifuged for 5 minutes at 13,000 rpm and the supernatant was transferred to a new test tube. The peptide was precipitated with trichloroacetic acid, the precipitate was washed with acetone and dissolved in water. TAMRA hydrazide (Lumiprobe, Russia) was added to the peptide solution to a concentration of 5 mM and incubated overnight at room temperature. After the reaction, the peptide was precipitated with trichloroacetic acid, the precipitate was washed with acetone and dissolved in water. Reverse-phase chromatography was performed on a Vydac C18 peptide column in an acetonitrile gradient with the addition of 0.1% trifluoroacetic acid to remove the unreacted dye.

### 4.4. Isolation of the nuclear fraction and preparation of chromatin-containing lysate from plant tissues

To isolate the nuclear fraction, 1 g of leaves (*Brassica rapa* subsp. pekinensis) were ground in a mortar with 3 ml of buffer A: 50 mM Tris-HCl, pH 7.4, 15 mM MgCl_2_, 10 mM KCl, 0.1% mercaptoethanol, 20% glycerol, 12.6 μl Protease Inhibitor Cocktail (Sigma, USA), and centrifuged for 10 min at 1260 g. The precipitate was re-suspended in 0.5 ml of buffer A with the addition of 3 % (v/v) Triton X100. Then 0.8 ml of 2.3 M sucrose was layered on the bottom of the centrifuge tube, and re-suspended precipitate was applied on the top and centrifuged for 10 min at 12000 g. Then the supernatant was removed, and the precipitate was washed twice with buffer A and centrifuged for 5 min at 12000 g. To obtain chromatin-containing lysate, the core precipitate was re-suspended in 1.5 ml of buffer B: 20 mM HEPES, pH 7.9, 1.5 mM MgCl_2_, 0.42 M NaCl, 0.2 mM EDTA, 25% (v/v) glycerol containing 1.5 μl 0.1 M DTT and 15 μl Protease Inhibitor Cocktail (Sigma, USA). This mixture was incubated with a gentle shaking for 30 minutes and, then, centrifuged at 20000 g for 5 min. The supernatant was transferred to a clean test tube and dialyzed against 0.1 M HEPES, pH 6.0. To remove H2A-H2B histones from the nucleosomes, a separate fraction of the supernatant was incubated for 10 min at 74^°^C. After incubation, the mixture was again centrifuged at 5 min at 20000 g. The precipitate was dissolved in 0.1 M HEPES, pH 6.0, and loss of histones H2A-H2B was checked by PAAG electrophoresis in comparison with the fraction not treated at 74^°^C.

### 4.5. In silico *prediction of potential association of the miPEP156a peptide with the cell nucleus and DNA*

Despite intensive research on the interactions between proteins and DNA over the past decades, the mechanisms of protein-DNA recognition remain poorly understood. Identification of nuclear proteins and potential DNA-binding regions of such proteins is an important step towards uncovering the functions of transcription factors. Today, various computational methods are widely used in this field to help identify nuclear proteins and predict possible protein binding sites. In this paper, we used fairly new hybrid algorithms to predict the nuclear localization of miPEP156a (http://mleg.cse.sc.edu/seqNLS/index.html) and the potential DNA-binding residues in this peptide [(https://dnabind.szialab.org/) and (http://mleg.cse.sc.edu/DNABind/)] (Liu and Hu, 2013).

### 4.6. Analysis of peptide binding to nuclear DNA and chromatin

The agarose gel shift method was used to analyze the binding of fluorescently labeled peptides to chromatin from Chinese cabbage and DNA (Rakitina et al., 2011). A non-specific DNA fragment of 280 base pairs long was a PCR product of the pUC18 plasmid. The peptide and DNA solutions were mixed in a buffer containing 10 mm HEPES pH 8.0, 50 mM KCl, 0.1 mM EDTA, 5% glycerol to obtain molar ratios of DNA/peptide from 1/25 to 1/200. The samples were incubated for 30 minutes at room temperature and mixed with a buffer for application to the gel. Electrophoresis was performed in 1.2% agarose gel. The presence or absence of 10 mM MgCl_2_ and mercaptoethanol (0.05% and 0.5%) did not affect the formation of the peptide – nucleic acid complex. In addition, to determine the stability of the peptide-DNA and peptide-chromatin complexes, total DNA from chromatin-containing lysate (size about 15,000 - 20,000 base pairs), or chromatin-containing lysate, were mixed together with a fluorescently labeled peptide in a ratio of 1μg peptide/10 ng DNA in a buffer containing 10 mM HEPES pH 8.0, 50 mM KCl, 0.1 mM EDTA, 5% glycerol. These mixtures were incubated for 10 min at room temperature and analyzed in 0.8% agarose gel.

### 4.7. Secondary structure of the miPEP156a peptide and its complexes with DNA

A solution of the miPEP156a peptide was prepared in water and as a mixture with total cabbage DNA (5 μg/ml), with a concentration of peptide - 0.012, 0.017 and 0.07 mM. The solutions were incubated at room temperature for 30 min. Then the circular dichroism (CD) spectra were measured on a J-810 spectropolarimeter (JASCO, Japan) in the range of 250 – 190 nm, in a quartz cuvette with a thickness of 0.05 or 0.1 cm at room temperature. CD spectra were recorded in increments of 0.2 nm (scanning speed 20 nm/min) and an optical slit width of 1 nm. The results were averaged over 4 spectra. To calculate the secondary structure, we used the CONTINLL program (CDPro package) and a set of SMP56 reference spectra.

### 4.8. Study of miPEP156a localization in plant tissue

4-days-old *B. rapa* roots were immersed in a peptide solution (23°C for 18 h) containing miPEP156a (10 μg/ml) labeled with the N-terminal TAMRA fluorophore in a MG buffer (10 mM MgCl_2_, pH 5.8). After treatment, the seedlings were washed three times with gentle shaking for 5 minutes in a MG buffer, treated for 60 minutes with a solution of the Hoechst 33342 fluorescent nuclear dye (0.625 mg/ml) (10 μl per 1.5 ml of buffer), and their roots and leaves were analyzed using confocal laser scanning microscopy (CLSM) at confocal microscope (Nicon Eclipse TE 2000-E, Japan). The fluorescence was visualized as follows: 495nm excitation/ 545nm emission.

### 4.9. Study of localization of the fluorescently labeled miPEP156a in animal cells

Assuming the affinity of miPEP156a for nucleic acids, we conducted experiments to study the intracellular localization of TAMRA-miPEP156a in mouse myeloma line Sp2/0 cells. The cells were seeded in DMEM medium with the addition of 10% FetalCS (Gibco, USA) in 24-well plates. The cells were seeded in such a concentration that the next day they were 40-60% of the monolayer. At the next day, TAMRA-miPEP156a (5 μg/ml) was added to the medium and incubated for 2 hours in a CO_2_ incubator. After 1 hour, 10 μl of the Hoechst 33342 nuclear marker (0.625 mg/ml) (Thermo Fisher Scientific, USA) was added, and the cells were incubated for another 1 hour. For the cell endomembrane labeling, 0.2 μl of the cytoplasmic marker ER-Tracker ™ Green (1 mM) (glibenclamide BODIPY ^®^ FL) (Thermo Fisher Scientific, USA) was added 50 min after the Hoechst 33342. Then the glasses with the cells were washed 3 times with a serum-free DMEM (Gibco, USA). After that, the glasses were removed from the tablet and attached to the slides with a glue. The intracellular localization of the labeled TAMRA-miPEP156a peptide, the ER-Tracker ™ Green cytoplasmic marker, and the Hoechst 33342 nuclear marker was studied using CLSM at confocal microscope (Nicon Eclipse TE 2000-E, Japan).

## 6. Acknowledgments

The authors thank Anastasia Ignatova (Shemyakin-Ovchinnikov Institute, RAS) for her assistance in experiments on measuring the secondary structure of the peptide using the circular dichroism method. This work was supported by a grant from the Russian Foundation for Basic research (grant No. 19-04-00174)

